# A contribution of FcγRIIIa cosignaling in T_FH_ subset development in Systemic Lupus Erythematosus

**DOI:** 10.1101/256198

**Authors:** Anil K Chauhan

## Abstract

**Background:** Expansion of follicular helper T cells (T_FH_) population occurs in systemic lupus erythematosus (SLE) and their numbers correlate with autoantibody titers. In this study, we sought to examine the role of ICs (Fc*γ*RIIIa costimulation) play in T_FH_ cells development.

**Methods:** We examined the presence of blood T_FH_ cells using multicolor flow analysis in SLE patients *in vivo*. We then examined the development of these cells *in vitro* using plate-bound ICs. Performed differential expression analysis in cells activated via Fc*γ*RIIIa and compared to CD28 cosignaling.

**Results:** In SLE patients PBMCs, CD4^+^ gated T cells show IC binding and phosphorylated spleen tyrosine kinase (pSyk). These pSyk^+^ cells express PD1, ICOS, IL-21, and Bcl6, the T_FH_ population markers. *In vitro* activation from plate-bound ICs of human naïve CD4^+^ T cells results in the differentiation of T_FH_ like cells phenotype. We show that Fc*γ*RIIIa-pSyk cosignaling in Bcl6^+^IL-21^+^ cells drives the production of both IFN-*γ* (T_FH_1) and IL-17A (T_FH_17) production. TLR9 engagement by CpG ODN 2006 combined with Fc*γ*RIIIa costimulation of CD4^+^ T cells augments, IL-17A, IL-21 production in Bcl6^+^ T cells. Fc*γ*RIIIa cosignaling induced the overexpression of microRNAs that participate in TLR signaling and are associated with T_FH_ cell differentiation. RNA-seq data reveal pathways that may contribute to the development of T_FH_ cells and nucleic acid sensing.

**Conclusion:** Our results suggest a role for Fc*γ*RIIIa receptors in T_FH_ development and a role for nucleic acid sensing in the expansion of T_FH_ cells.

## Introduction

Systemic lupus erythematosus (SLE) is associated with elevated levels of immune complexes (ICs) formed with antibodies against altered self-nucleic acids. Nucleic acid containing ICs activate innate immune pathways and nucleic acid sensing in innate cells triggers interferon (IFN) responses. Such events have not been explored in CD4^+^ T cells. Increased numbers of circulating T_FH_ cells are present in SLE patients. T_FH_ cells are being evaluated as therapeutic target for SLE [1, 2]. T_FH_-like cells that express Bcl6, PD1, ICOS and produce IL-21 are present both in ectopic germinal centers (GCs) and lymphoid aggregates [3]. Three subsets of T_FH_ cells are recognized and expression of Bcl6 in circulating T_FH_ cells is debatable [4–6].

Ectopic GCs are observed in nephritic kidney in SLE, which show abundance of ICs and complement deposits. The evidence that ICs may play a critical role in GC reaction comes from many previous studies. In GCs, FDCs, macrophages and B cells retain ICs on the cell surface and fulfils the constant antigenic requirement for T_FH_ cell development [7]. Subcapsular sinus (SCS) macrophages retain intact ICs on the cell surface, and transfer these intact ICs to the follicular B cells. Cognate B cells upon capturing ICs migrate to the T cells zones [8, 9]. B cells then scan FDC processes for ICs and upon acquiring ICs from FDCs, they make a contact with T_FH_ cells before moving to GC reaction phase [10]. B cells capture ICs from the macrophages via complement receptors (CRs) and facilitate their transport to FDCs [9]. These studies suggest a key role for ICs in the GC reaction. A role for CRs in the proposed IC transfers is suggested, however a role for Fc receptors (FcRs) in these IC transfers is not excluded. In tightly packed lymphoid follicles, ICs transfer occur from a non-cognate antigen transporting B cells to antigen specific cognate B cells [9]. Furthermore, a non-cognate transport of ICs from B cells to FDCs also occurs [11]. A role for Fc*γ*RIIIa costimulation in naïve/memory cells and ICs bound to T_FH_ cells surface in the GCs reaction is important area of investigation for the better understanding of autoimmune pathology. T_FH_ cells upon reaching GCs, trigger T_FH_-GC B cell interactions, which establishes serological memory [10, 12]. Costimulatory proteins i.e. CD40L-CD40, ICOS-ICOSL, accessory molecules OX40-OX40L, as well as cytokines and their receptors in particular IL4-IL4R, IFN-*γ*-IFN-*γ*R, IL21-IL21R and SAP-SLAM are protein pairs that contribute to the T_FH_-GC B cell interactions [12]. The disruption of these interaction in SLE mouse models ameliorate disease.

We report that the circulating T_FH_ like cells in blood express Fc*γ*RIIIa and show pSyk. In the SLE peripheral blood mononuclear cells (PBMCs), this subset of CD4^+^ T cells express T_FH_ markers i.e. Bcl6, ICOS, PD1, IL-21, CXCR3 and CXCR5. Additionally, we show that the *in vitro* costimulation of Fc*γ*RIIIa from the engagement of plate-bound ICs in the presence of sub optimal anti-CD3 ligation, drive the generation of Bcl6^+^ T_FH-_like cells that produce IFN-g and IL-17A. In addition to Fc*γ*RIIIa-Syk cosignaling, TLR9 ligation by CpG ODN 2006 in human naïve CD4^+^ T cells enhance IL-21 and IL-17A production in Bcl6^+^ cells. Fc*γ*RIIIa cosignal in naïve CD4^+^ T cells upregulate microRNAs (miRNAs) that contribute to T_FH_ differentiation and TLR signaling.

## Materials and Methods

### ICs and C5b-9

ICs were purified from 50 ml of pooled serum or plasma from five to ten SLE patients with high levels of complement opsonized ICs. The purification procedures for ICs and C5b-9 have been previously described [13–15]. The nature of the ICs was characterized for binding to Fc*γ*R using cells that express this receptor and compared with aggregated human *γ*-globulin (AHG) and anti-Fc*γ*RIIIa antibody (clone 3G8) [16]. In addition, ICs were tested for their potential to activate CD4^+^ T-cells and compared with *in vitro* formed Ova-anti-Ova ICs in an earlier study [13].

### T cell isolation and Activation

Blood was collected with informed consent in the Saint Louis University Rheumatology clinic. The PBMCs were isolated using Histopaque gradient (Sigma). PBMCs were isolated within 12 h of sample collection and monocytes were removed by overnight plating in a culture dish. The next day, the CD4^+^CD45RA^+^ cells were purified using naïve CD4^+^ T-cell isolation kit II (product no. 130-094-131, Miltenyi Biotec). Purified naïve CD4^+^ T-cells represented a >97% pure population. Purified cells were maintained in culture with 20 U of IL-2 for two days. Thereafter, these cells were stimulated with plate-bound ICs at 10 μg/ml opsonized with soluble C5b-9 at 2.5 μg/ml for each 2.5 × 10^5^ cell in the presence of plate-bound anti-CD3 at 0.25 μg/ml (eBioscience, clone OKT3). Positive controls cells received 1 μg/ml of anti-CD28 (BD biosciences, clone 28.2). All studies were performed in the presence of anti-CD3, unless specified. Cells were polarized in the presence of IL-1β, IL-6, TGF1-β, and IL-23 as reported earlier [17]. For inhibition experiments, cells were treated with 10nM of P505 for sixty minutes, a Syk inhibitor (product no. PRT06207, Selleckchem) prior to activation and were cultured in the presence of P505. Cells were activated with plate-bound ICs for 48 h in the presence of IL-2 (20 IU/ml) medium (Peprotech). Post 48 h, cells were again re-stimulated and processed for staining at 96 h.

### Flow cytometry analysis

Multicolor flow staining was performed immediately after purifying the PBMCs. Cells were stained using anti-human CD4 PE-Cy7 (RPA-T4), anti-human CD4 PE eFluor 610 (RPA-T4), anti-human pSyk-PE (moch1α), anti-human ICOS-APC eFluor780 (1SA-3), anti-human-PD1-PE-Cy7 (eBio J105), anti-human CXCR5 eFluor 450 (MUSBEE), all from eBiosciences. Other antibodies used anti-human Bcl6-PE-CF594 (K112-91, anti-human IL-17A-APC-R700 (N49-553), anti-IL-21-BV421 (3A3-N2.1), anti-human IFN-*γ*-BV711 (B27), anti-human CXCR3-BV711 (1C6/CXCR3) were purchased from BD Biosciences. Another anti-human IFN-*γ*-APC (4S.B3) was purchased from Biolegend. ICs were labeled using Alexa Fluor 488 5-TFP ester (A3005) from Molecular Probes. The ICs conjugates were in the range of 23 to 30 μM fluorochrome to-protein ratio. Purification and conjugation was performed as described previously [16, 18]. All conjugates were titrated and flow compensation was done using isotype controls. Cell surface staining was performed at RT for 30 minutes, thereafter cells were washed and fixed using cytofix buffer for 30 minutes. Cells were washed again and intracellular staining for cytokines and transcription factor was performed in permeabilization buffer for 60 minutes. Both buffers were purchased from eBiosciences. Flow staining for PBMCs were carried out using multicolor flow analysis using various antibody conjugates in different panels such as CD4, ICs, Bcl6, pSyk, IL-17A, IFN-y; CD4, ICs, ICOS, PD1, pSyk, IFN-*γ*, CXCR3, CXCR5; CD4, ICs, PD1, ICOS, Bcl6, IL-21, CD4, ICs, IFN-*γ*, IL-17A, IL-21, Bcl6, pSyk. In the PBMCs, FSC, CD4^+^ T cells were first gated and analyzed for the expression of various markers. In all analysis performed i*n vitro* experiments CD4^+^ T cells were analyzed unless specified in the figure. A two-tailed paired non-parametric t-test and correlation coefficient was calculated using Prizm software unless specified.

### CpG ODN 2006 treatment of CD4^+^ T cells

For studying the effect of CpG ODN 2006, cells were stimulated and polarized in the presence of IL-1β, IL-6, IL-23, and TGF-β as described earlier [17]. Purified naïve CD4^+^ T-cells at a density of 2.5 × 10^6^ were plated in 96-well plates (Nunc) and activated [16, 17]. Colonies were broken at every 48 h of culture. On day seven CpG and non-CpG ODN at a final concentration of 5 μM/ml were added. This concentration was used since human PBMCs and CD4^+^ T-cells respond at this concentration for cytokine production [19, 20]. ODN 2006 TC*GTC*GTTTTGT C*GTTTTGTC*GTT (CpG ODN) and control ODN TGCTGCTTTTGTGCTTTTTGTGCTT were synthesized by IDT (USA). We analyzed these cells for IL-17A and IL-21 production without PMA and ionomycin stimulation.

### RNA-seq transcriptome analysis

Purified were differentiated using a mixture of four cytokines as described in previous section. RNA was purified from naïve CD4^+^ T-cells activated with anti-CD3 (0.25 μg/ml) + anti-CD28 (2.0 μg/ml) or anti-CD3 + ICs opsonized with C5b-9 using an RNA purification kit from Clontech laboratories. A total of 20 ng of RNA was used to generate the RNA-seq libraries using the Ion AmpliSeq Transcriptome Human Gene Expression Kit as per manufacturers recommendations. The libraries were indexed using the primers from ABI. We then sequenced these synthesized libraries using the Ion Torrent Sequencer in the Genomic Core (Saint Louis University School of Medicine). Statistical analysis and differential expression was performed for RNA transcripts using R and subjected to GO annotation analysis using R. Heat maps and cluster analysis was performed using CIMinor (nci.nih.gov). Reads per million from the RNA-seq data was used as input for the generation of the heat maps. Total normalized reads obtained from Fc*γ*RIIIa or CD28 cosignaling was divided with reads obtained from untreated paired sample of the same subject. Relative fold change was used for generating heat maps. Gene expression dot plots were generated in R.

## Results

### In SLE, ICs bound cells express T_FH_ markers and show phopshorylated Syk

T_FH_ cells participate in the pathogenesis of primary immunodeficienies, autoimmunity, inflammation and cancer [1]. In circulation three subsets of blood memory follicular cells i.e. T_FH_1, T_FH_2, and T_FH_17 are recognized based on the expression of cytokines and chemokines receptors [1]. Upon binding of ICs, Fc*γ*RIIIa signals by phopshorylating Syk. To determine whether Fc*γ*RIIIa costimulation contributes to the T_FH_ development, we examined the binding of ICs, and pSyk in CD4 gated population that express T_FH_ markers. A subset of CD4^+^ T cells in the SLE PBMCs bound to ICs and these cells showed pSyk (figure 1a, b). We then analyzed whether these cells express Bcl6, a transcription regulator of T_FH_ cells. Results showed that the IC bound and pSyk^+^ cells expressed Bcl6 (figure 1c, d and figure S1). These results for Bcl6 expression are in agreement with two previous studies from SLE subjects [6, 21]. Further analaysis showed that a subset of the Bcl6^+^ cells also produced IFN-*γ* (figure 1e). pSyk^+^Bcl6^+^ cells showed strong correaltion with IFN-*γ*^+^Bcl6^+^population (r = 0.948) suggesting a role for Syk activation in IFN-*γ* production [16]. A strong correlation among pSyk^+^Bcl6^+^ and ICs^+^Bcl6^+^ as well as ICs^+^Bcl6^+^ and IFN-*γ*^+^Bcl6^+^ cells was observed (figure 1f). These findings are consistent with our earlier report, where we observed IFN-*γ* production from Fc*γ*RIIIa cosignaling in activated human naïve CD4^+^ T cells [16, 17]. Blood memory T_FH_1 cells are marked by the presence of chemokine receptor CXCR3^+^ and IFN-*γ* production [22]. Circulating T_FH_1 cells correlate with the generaton of protective antibody repertoire [22, 23]. To examine the participation of Fc*γ*RIIIa cosignaling in IFN-*γ* production in T_FH_1 cells, we analyzed CXCR3 expression and IFN-*γ* production in pSyk^+^ cells. These results indicated that the CXCR3^+^IFN-*γ*^+^ double positive cells show strong correlation with the CXCXR3^+^pSyk^+^ cells, r = 0.910 (figure 1i). These cells likely represent memory effector population, since lack of CXCR3 expression marks T central memory cells [5]. Post influemza vaccination CXCR3^+^ T_FH_ cells were able to induce memory B cells to differentiate into antibody producing plasma cells [23]. Changes in the frequency of T_FH_1 cells show association with disease activity [24]. In two patients, we observed cells that produced CXCR3 but did not show pSyk and produced IFN-*γ* (figure S2). These results, thus suggest that for Fc*γ*RIIIa-pSyk cosignaling possibly does not contribute to the CXCR3 expression observed in blood T_FH_1 like cells.

**Figure 1.**
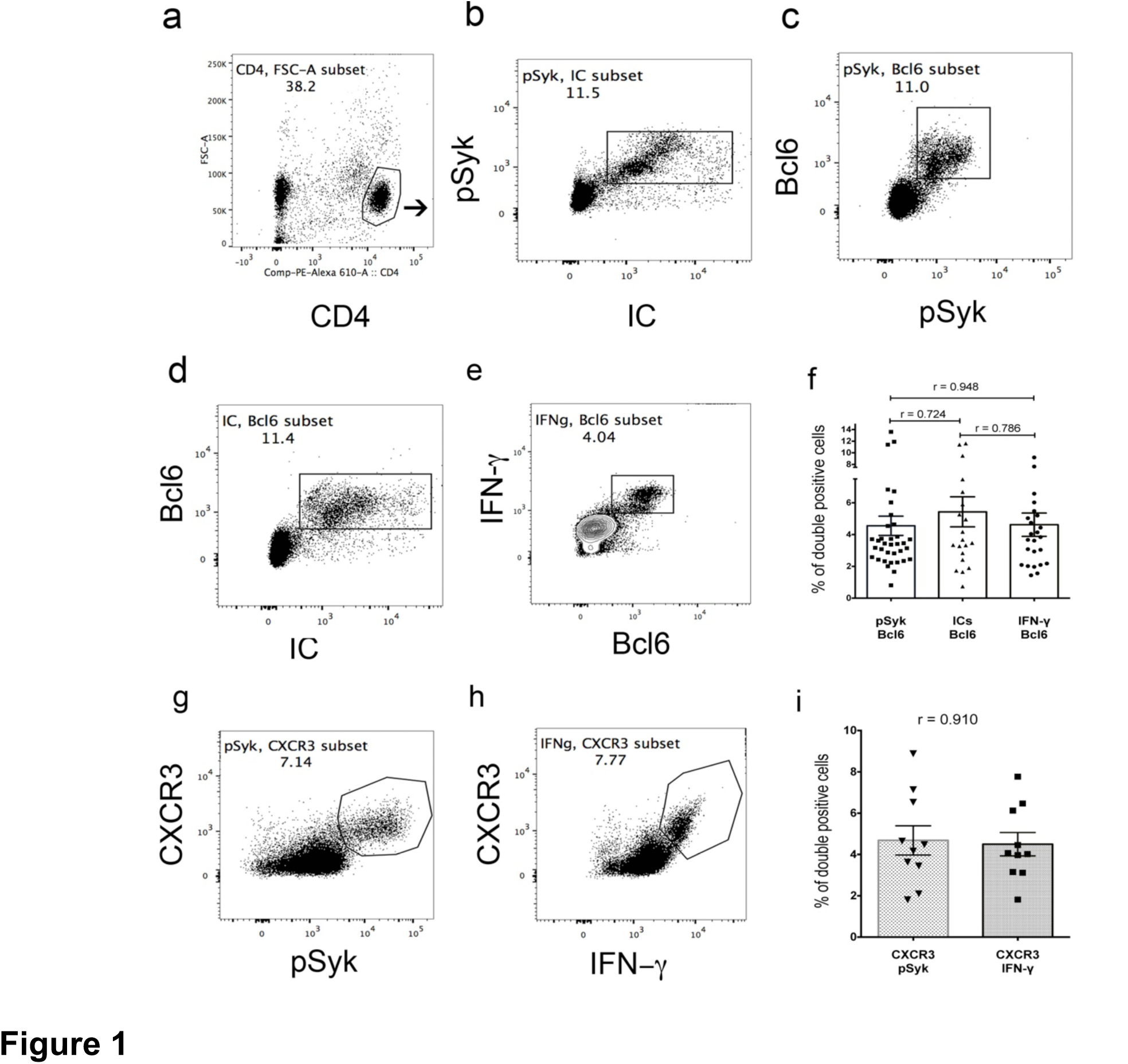
In SLE PBMCs, CD4^+^Fc*γ*RIIIa^+^ T cells show T_FH_ markers. Flow cytometry analysis of CD4^+^ gated T cells (a) show cell population that bound to IC and show pSyk^+^ (b). IC^+^ and pSyk^+^ cells express Bcl6 (c, d). A subset of Bcl6^+^ cell produces IFN-*γ* (e). pSyk^+^Bcl6^+^: ICs^+^Bcl6^+^ (n = 21, r = 0.724); IFN-*γ*^+^Bcl6^+^: pSyk^+^Bcl6^+^ (n =25, r = 0.948) and IC^+^Bcl6^+^: IFN-*γ*^+^Bcl6^+^ (n= 21, r = 0.0786) cells show strong correlation (f). pSyk^+^ cells also express CXCR3 (g) and produce IFN-*γ* (h). CXCR3^+^ cells also show a strong correlation among pSyk^+^ and IFN-*γ*^+^ production (n = 10; r = 0.910).

To further establish that the Bcl6^+^pSyk^+^ cells represent T_FH_ like subset, we next examined the expression of ICOS, CXCR5, and PD1 in a multicolor flow analysis. This analysis showed that pSyk^+^ cells expressed ICOS, CXCR5 and PD1 (figure 2a, b). These results thus suggest that the Fc*γ*RIIIa-pSyk signaling contribute to circulating T_FH_ cells. PD1 expression define acitvated and quiescent memory T_FH_ cells and correalte with disease activity [5]. A higher percentage of cells that lacked pSyk showed higher PD1 expresion (PD1^high^) contrary to pSyk^+^ cells that showed moderate amount of PD1 (PD1^mod^) at a *p Value* of <0.023 (figure 2d). pSyk^-^ cells showed statistically signficant higher MFI for PD1 at a *p Value* of <0.001 compared to PD1 in pSyk^+^ cells. Src homology 2 domain-containing protein tryosine phopshatase (SHP-2) binds to PD1 and PD1-SHP2 complexes dephophorylates CD28 effciently in intact cells [25]. Such complexes may contribute to the partial Syk dephosphorylation in PD1^high^ population resulting in the development of PD1^mod^ cells. In our previous publication, we have shown that pSyk^+^ cells represent activated cells that also express CD25, CD69 and CD98[17]. Another low affinity FcR, Fc*γ*RIIa is expressed by CD4^+^ T cells, which also express PD1 and these cells are carrier of HIV provirus [5]. Our data suggests that Fc*γ*RIIIa-pSyk signaling has a role in the expansion of circulating blood T_FH_ cells.

**Figure 2.**
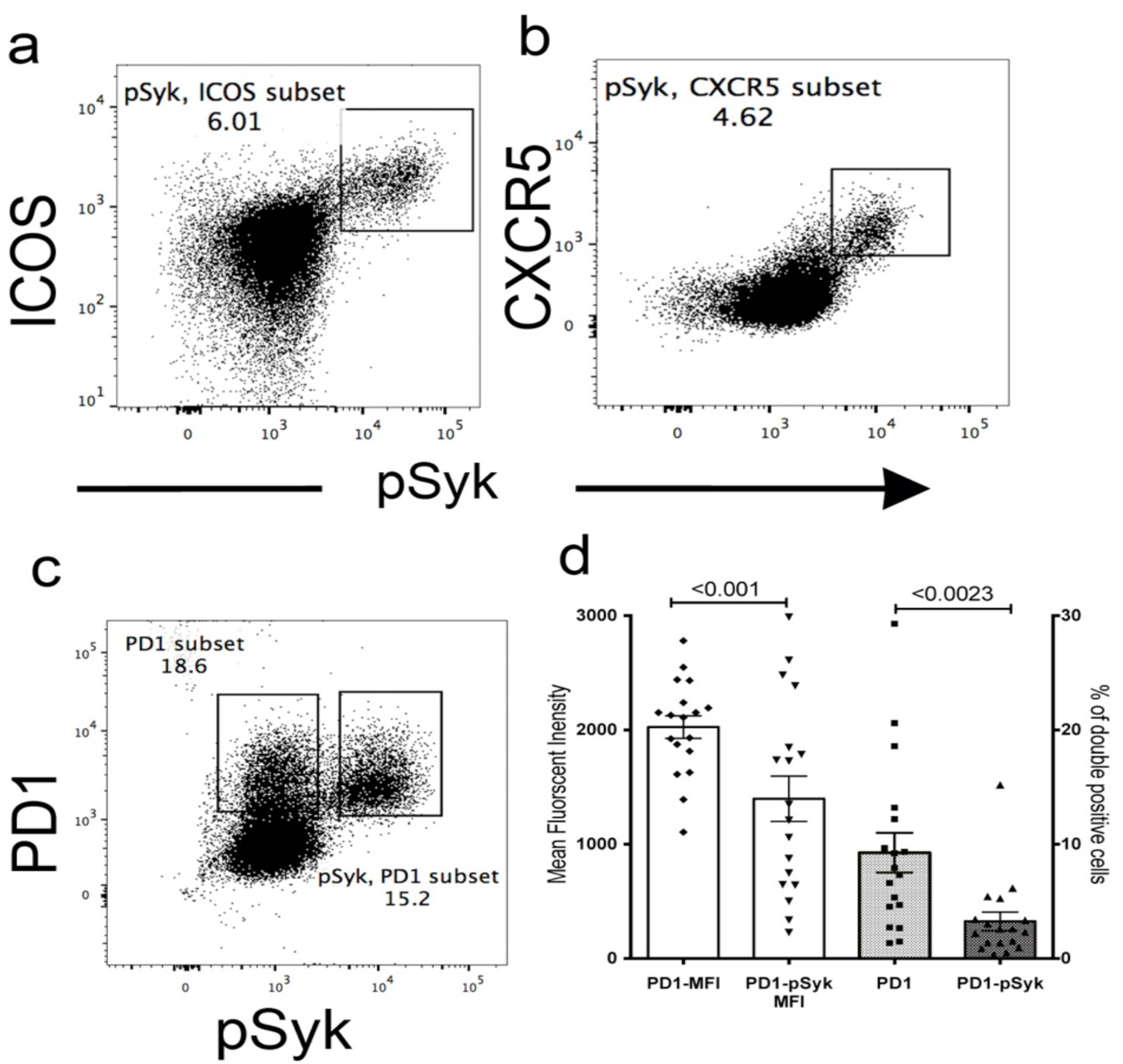
In SLE PBMCs, pSyk^+^CD4^+^ T cells express ICOS, CXCR5 and PD1. CD4^+^pSyk^+^ T cells express ICOS (a) and CXCR5 (b). Two populations of PD1 expressing cells PD1^high^ pSyk^-^ and PD1^mod^pSyk^+^ are observed (c). The percentage of PD1^mod^pSyk^+^ cells is statistically significant lower than PD1^high^pSyk^-^ (*p value* = <0.001) and they also show lower amount of PD1 (MFI values) (*p value* = <0.0023) (d). Shown representative flow plot of sample analyzed in four experiments.

### *In vitro* Fc*γ*RIIIa engagement by ICs induces T_FH_ like cells

To further confirm a role for Fc*γ*RIIIa-pSyk signaling in the generation of T_FH_ cells, we examined the expression of T_FH_ markers in human naïve CD4^+^ T cells that were activated *in vitro* by using plate-bound ICs in the presence of suboptimal anti-CD3. This activation resulted in ICOS and PD1 expression in pSyk^+^ cells (figures 3Aa, b, c, d and e). These data thus suggest that Syk activation participate in T_FH_ devlopment. A small population of pSyk^-^ cells did not express ICOS. Unlike *in vivo* data, all pSyk^+^ cells expressed PD1. We have previously shown and compared the IFN-*γ* and IL-17A production in the condition used here from both Fc*γ*RIIIa and CD28 costimulation [17]. A similar results were also obtained for IL-21 and Bcl6 expression (figure S3). Upon *in vitro* Fc*γ*RIIIa-pSyk cosignaling, we did not observe the PD1^high^pSyk^-^ population and all cells that were PD1^+^ also showed pSyk. These results thus suggest the possible dephosphorylation of pSyk in PD1^high^ cells in circulation by a serum phosphatase that associates with PD1. Alternatley, these cells lacked the induced expression of Fc*γ*RIIIa. IL-21 is a signature T_FH_ cytokine that was upregulated by Fc*γ*RIIIa-pSyk cosignaling. IL-21 blocks follicular regulatory (T_FR_) response and drive B cell activation and differentiation [26, 27]. During autoimmune response, Fc*γ*RIIIa mediated excessive IL-21 production could overcome T_FH_ suppression by the regulatory effect of T_FR_ cells. We also examined IL-21 production in Bcl6^+^ cells from Fc*γ*RIIIa-pSyk cosignaling. Bcl6^+^ cells produced IL-21 (figure 3Af). A subset of the Bcl6^+^ cells produced IFN-*γ* (figure 3Ag). We further confirmed the expression of Bcl6 as RNA transcripts by RT-PCR in cells that were activated via Fc*γ*RIIIa cosignaling normalized against untreated control cells (figure 3Ah). These results showed upreguation of Bcl6 RNA transpcripts. Bcl6 RNA transcripts have been reported in circulating T_FH_ cells [22]. We next examined, whether Bcl6^+^pSyk^+^ positive cells that produce IL-21 also produce IFN-*γ* or IL-17A (figure 3Ba, b, c). Results showed that a subset of Bcl6^+^pSyk^+^ cells produced IL-21^+^ and also produced IFN-*γ* or IL-17A. IFN-*γ* production was observed in higher percentage of cells compared to those that produced IL-17A, which was statistically significant at a *p Value* of <0.0001 (figure 3Bd). These data suggest that the Fc*γ*RIIIa-pSyk cosignaling likely participate in the development of both T_FH_1 and T_FH_17 populations. This raises the possibility that in autoimmunity, pathogenic ICs contribute to the differentiation of both T_FH_1 and T_FH_17 cells. To further confirm that the Syk mediated signaling participate in IL-21 production, we tested the effect of a Syk inhibitor P505 on IL-21 production and Syk phosphorylation. At 10 nM concentration, P505 successfully blocked Syk phosphorylation and inhibited IL-21 production (figure 3Be, f). This data thus suggest a role for Fc*γ*RIIIa-pSyk cosignaling in the differentiation of blood circulating T_FH_ subsets.

**Figure 3.**
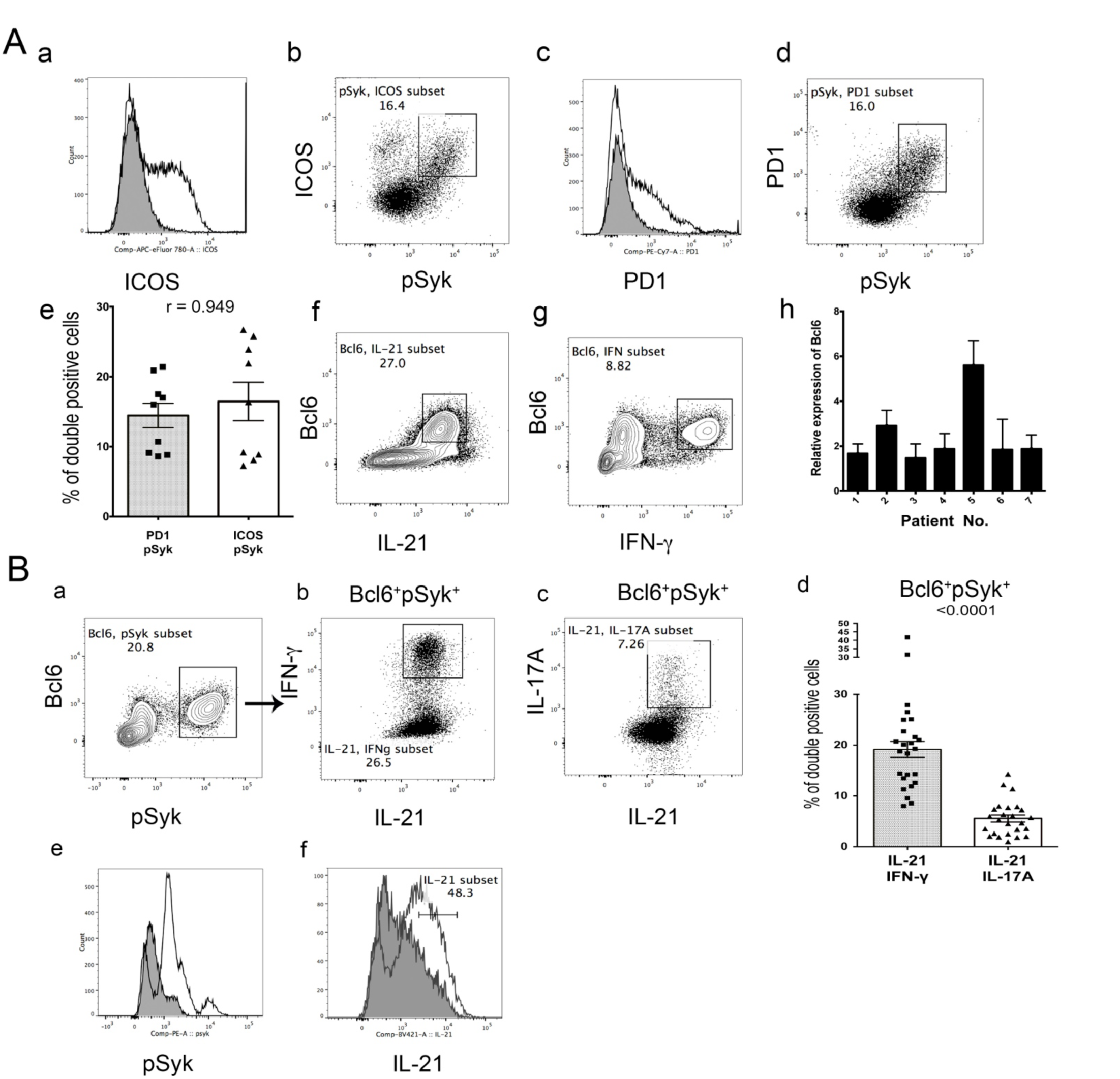
Purified naïve CD4^+^ T cells activated *in vitro* with plate-bound ICs express T_FH_ markers. **(A)** IC treated cells express ICOS in pSyk^+^ cells (a, b). These cells also express PD1 in pSyk^+^ cells (c, d). A strong correlation between ICOS and PD1 expression is observed, r = 0.949, n =9 (e). IC treatment induces Bcl6 expression and IL-21 production (f). CD4^+^ T cells in PBMCs also show cells that express Bcl6 and produce IFN-*γ* (g). Bcl6 transcripts are observed in IC treated cells, normalized on the levels of expression in paired untreated cells (h). **(B)** Bcl6^+^pSyk^+^IL-21^+^ cells produce IFN-*γ* (b) and IL-17A (c). IFN-*γ*^+^ is produced by higher percentage of cells compared to IL-17A, n = 25 (*p Value* of <0.0001) (c). P505 treated cells (shaded) reduced pSyk^+^ population compared to IC treated control cells (open) (e). P505 also inhibited IL-21 production (shaded) compared to ICs treated control cells (open) (f). Shown 1 of 4 analyzed in two experiments.

### TLR9 signaling synergize with Fc*γ*RIIIa costimulation to enhance IL-21^+^Bcl6^+^ and IL-17A^+^Bcl6^+^ populations

Up-regulation of TLR9 signaling in monocyte-derived DCs promote enhanced T_FH_ differentiation [28]. In human B cells, cosynergistic signaling from ITAM and MyD88 enahnces B cell receptor activation [29]. We previously reported that the TLR9 and Fc*γ*RIIIa joint cosignaling enhances IL-17A and IL-21 production [18]. Thus, we analyzed IL-17A and IL-21 prodution from synergistic cosignaling in Bcl6^+^ T_FH_ cells. We compared the percentage increase in Bcl6^+^ cells that produced IL-21 and IL-17A. Results showed that the joint signaling by Fc*γ*RIIIa-pSyk and TLR9 significantly enahnced the population of double positive Bcl6^+^IL-21^+^ (*p Value* <0.0001) and Bcl6^+^IL-17A^+^ (*p Value* <0.0003) populations compared to control cells that were only treated with CpG ODN 2006 alone (figures 4 A-C). We also examined the diffrences in these populations upon ODN (nonphosphothiorated) and CpG ODN treatments. A statistically significant increase in Bcl6^+^IL-21^+^ producing cells was observed from ICs + CpG ODN treatment, compared to control group (*p Value* <0.0001). An increase in this population was also observed from ODN treatment compared to control cells (*p Value* <0.0005). This analysis showed a statistically significant increase from phosphothiorate modification of ODN (CpG), compared to ODN in combination with the ICs treatment (*p Value* <0.0145). Similar results were observed for Bcl6^+^IL17A^+^ population. Statistically significant increase in Bcl6^+^IL-17A^+^ cells were also observed upon ICs + CpG ODN treatment compared to control cells (*p Value* <0.0003). Bcl6^+^IL-17A^+^ cells showed significant increase between CpG ODN vs. ODN (*p Value* <0.043). Both naïve CD4^+^ T cells obtained from SLE or normal subjects showed these changes and there was no observed difference between these groups. It is known that TLR9 recognize both phosphothiorated modified and unmodified ODNs [30]. Our results confirm these previous findings in the generation of T_FH_17 like cells. Our data suggest that the Fc*γ*RIIIa-pSyk cosignaling synergize with TLR9 to enhances the development of T_FH_ like cells. Both CpG modified ODN and unmodified ODN drive these changes. In the presence of ODN, PMA and ionomycin stimulation disproportionately affects IL-21 production, thus we performed studies in the absence of these stimulants. This affect of PMA and ionomycin is in agreement with similar effect of these agents on T_FH_ cell generation in previous studies [1].

**Figure 4.**
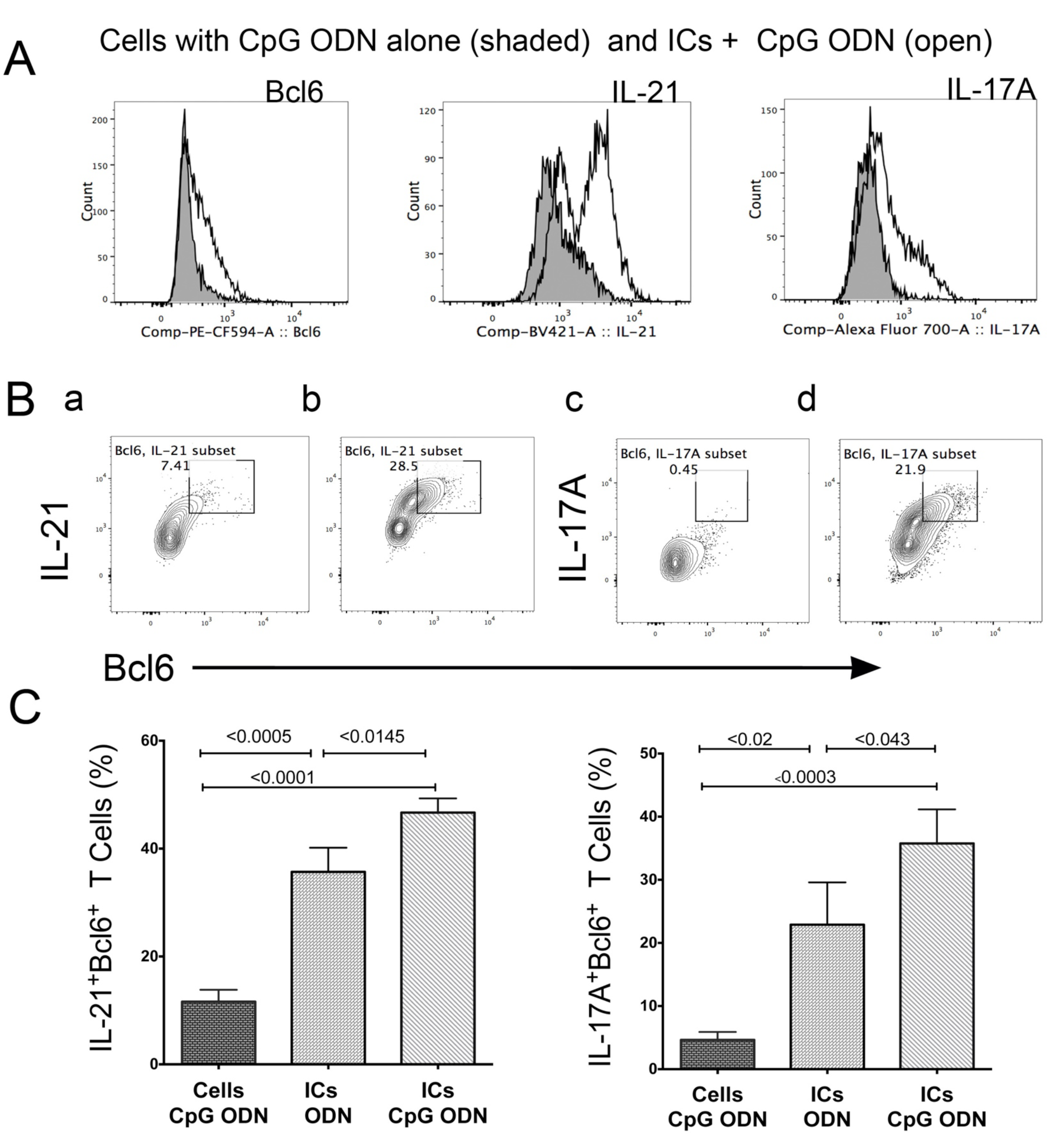
In purified naïve CD4^+^ T cells, CpG 2006 co treatment with ICs enhances IL21 and IL-17A production in Bcl6^+^ cells. **(A)** IC engagement of Fc*γ*RIIIa and ODN CpG 2006 with TLR9 enhances Bcl6 expression (a), IL-21 (b) and IL-17A production (c). Control cells treated with CpG ODN2006 (shaded). **(B)** Bcl6^+^ cells show enhanced IL-21 production from joint signaling (a) compared to control treated with ODN CpG 2006 (b). A similar result is observed for IL-17A, joint signaling (c) and control (d). **(C)** Statistically significant increases are observed with both ODN and CpG ODN treatment compared to control cells for the expression of IL-21^+^Bcl6^+^ (a) and IL-17A^+^Bcl6^+^ cells (b). *p Values* shown above the corresponding bars (n=8). All treatments were performed in the presence of suboptimal anti-CD3.

### Fc*γ*RIIIa is a distinct costimulation than CD28 cosignal

RNA-seq analysis data suggest that the Fc*γ*RIIIa cosignaling upregulate different signaling pathways compared to CD28 cosignaling. Combined analysis from four paired samples from Fc*γ*RIIIa costimulation showed upregulation of 1407 gene transcripts vs. 402 from CD28 cosignaling (figure 5). These data suggest that Fc*γ*RIIIa is a distinct costimulatory signal in CD4^+^ T-cells. Fc*γ*RIIIa activation represents a state of stress, since a number of genes that were upregulated are stress response genes. A number of heat shock proteins were upregulated upon Fc*γ*RIIIa cosignaling [18]. A significant increase in the expression of G-protein coupled receptor (GPCR) pathway proteins was observed upon Fc*γ*RIIIa costimulation. A 4.9-fold enrichment of 204 genes from GOTERM_PBGO:0007186, GPCR proteins signaling pathway at a FDR of 7.98E-89 and at a *p Value* of 4.71E-92 was observed. Interestingly, 223 genes from BPGO:0007166, cell surface receptor linked signal transduction category showed 3.2-fold enrichment at a FDR value of 1.72E-62 at a *p Value* of 1.02E-65. The GPCR signaling proteins regulate the expression of several chemokine’s and their receptors. These proteins are critical in homing of the T_FH_ cells into tissue and their exit into circulation.

**Figure 5.**
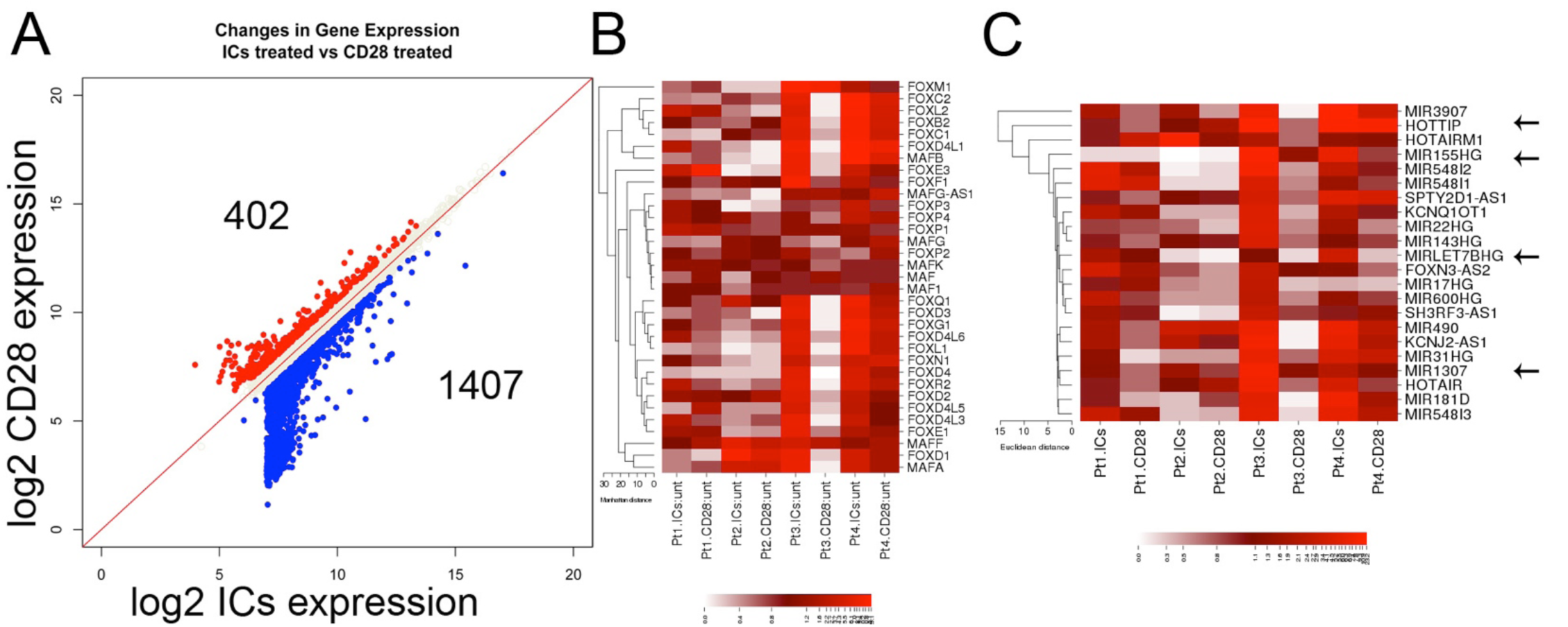
Differential gene expression Fc*γ*RIIIa vs. CD28 cosignaling. Dot plot showing differential gene expression from CD28 and Fc*γ*RIIIa cosignaling pathways **(A)**. Cluster analysis and heat maps show induced expression for FOXO and c-MAF transcription factors upon Fc*γ*RIIIa cosignaling that are associated with T_FH_ development **(B)**. Heat map showing cluster analysis of microRNAs expressed from Fc*γ*RIIIa and CD28 cosignaling in four patients **(C)**. MIR1307 is a risk allele for SLE pathogenesis. Patients 1, 3, and 4 show strong upregulation of key microRNAs (marked with arrow). Color scale 0 to 23.1 (B), and 0 to 23.2 (C).

We next examined the expression of transcription factors c-MAF and FOXO that regulate T_FH_ responses. Analysis showed that RNA transcripts from both of these pathways were upregulated upon Fc*γ*RIIIa costimulation compared to CD28 (figure 5B). ICOS signaling activates PI3K-Akt signaling, a GPCR mediated pathway, which also regulate FOXO1 ubiquitination. A large set of genes from ubiquitination pathways are upregulated upon Fc*γ*RIIIa costimulation in CD4^+^ T cells (AKC, unpublished observation). These data further support our hypothesis that Fc*γ*RIIIa-pSyk cosignaling contributes to the T_FH_ cell expansion

### Fc*γ*RIIIa cosignaling upregulates miRNAs that are associated with T_FH_ development

MicroRNAs regulate CD4^+^ T cell differentiation [31, 32]. miR-155, miR-146a, as well as miR-17~92 cluster (miR-17 is part of this cluster) regulate the development of T_FH_ cells [4, 33]. To examine the participation of miRNAs, we examined and compared the levels of miRNAs RNA-transcripts from Fc*γ*RIIIa costimulation and compared them with CD28 costimulation. Hierarchical cluster analysis showed upregulation of several key miRNAs from Fc*γ*RIIIa costimulation, such as miR-155 (MIR155HG) (host gene, HG), miR-Let 7B (MIRLET7BHG), miR-17HG, and miR-3907. These RNAs were prominently overexpressed in three of four patients analyzed (figure 5C). miR-155 was upregulated in two of four subjects analyzed. MiR-155 regulates IRAK3, and IKKe, which then via IRF3 induces type I IFNs [34]. miR-155 also promotes T_FH_ cells accumulation during chronic low-grade inflammation [35]. A large subset of miRNAs and lincRNAs were modulated by Fc*γ*RIIIa costimulation vs. CD28 in naïve CD4^+^ T cells (figure S4). miR-3907, which is a known risk allele (GWAS study) was upregulated upon Fc*γ*RIIIa costimulation (figure 5C) [36]. miR-22 by influencing HDAC4 regulate T cell differentiation. miR-490 modulates cell growth and miR-548 targets IFNgR1. miR-548I1-3 associates with RNA-induced silencing complex (RISC) and regulates translation imitation. These cumulative data suggest that the Fc*γ*RIIIa cosignaling via miRNAs, regulates the T_FH_ differentiation. Both HOTTIP and HOTAIR were also upregulated. These miRNAs are shown to regulate several key autoimmune processes.

Our data from both direc*t in vivo* and *in vitro* analysis provide a strong evidence for the participation of Fc*γ*RIIIa cosignaling the generation of T_FH_ like cells, which will form cytoconjugates with B cells and trigger the generation of autoreactive B cells (figure 6).

**Figure 6.**
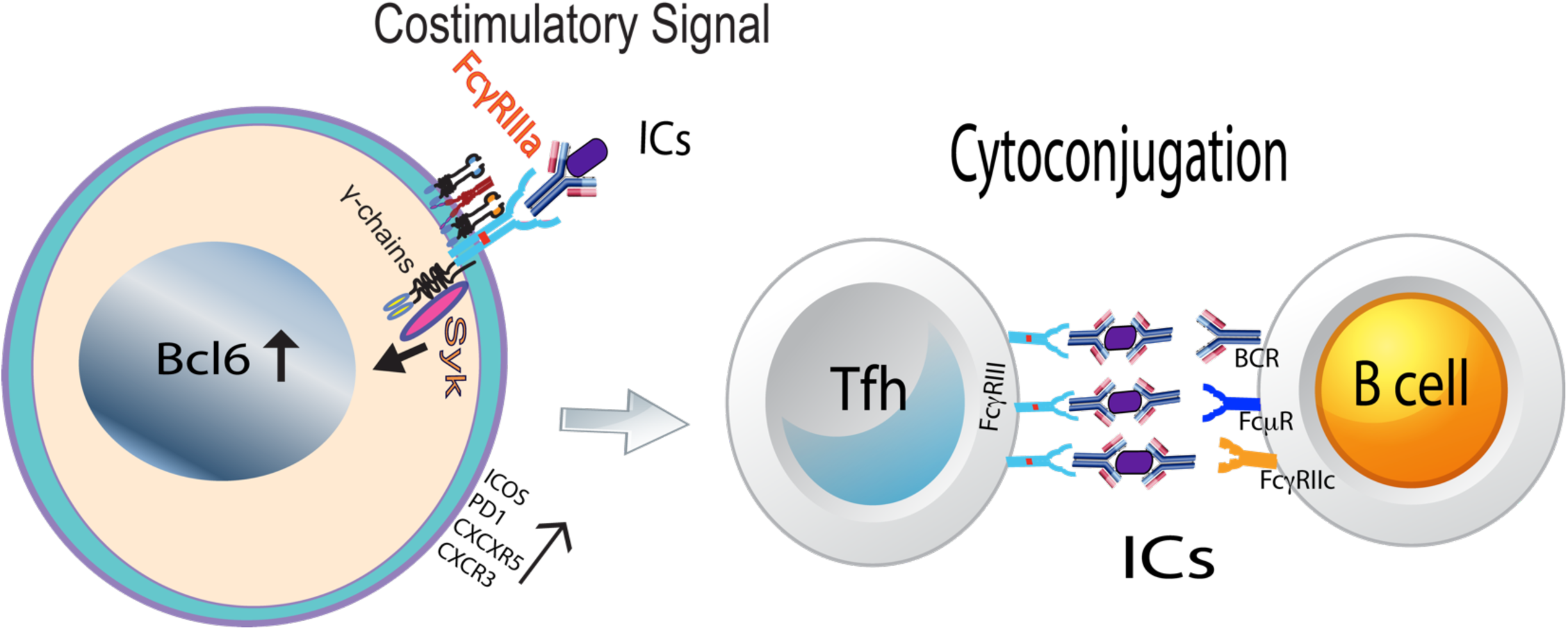
Proposed model of the role for Fc*γ*RIIIa cosignaling in T_FH_ responses. Fc*γ*RIIIa cosignaling from the ligation by ICs in CD4^+^ T cells triggers Syk phosphorylation. Syk signaling contributes to the generation of T_FH_ cells. T_FH_ cells, which express Fc*γ*RIIIa engage ICs and would facilitate the cytoconjugation with B cells. On B cells ICs could simultaneously engage BCR, FcμR or Fc*γ*RIIc.

## Discussion

Circulating blood T_FH_ cells are increased in SLE and their number correlate with disease activity [24, 37]. Thus, it is important to identify the signals that contribute to their expansion and characterize their phenotype in SLE [12]. In this study, we show that the Fc*γ*RIIIa-pSyk cosignaling, which is a positive costimulatory signal for naïve CD4^+^ T cells, contributes to the T_FH_ cells generation. In SLE PBMCs, CD4^+^ T cells express Bcl6, produce IL-21, bind to ICs and are pSyk^+^. Second *in vitro* activation of Fc*γ*RIIIa-pSyk pathway from plate-bound ICs generate T_FH_ like cells, suggesting a role for Fc*γ*RIIIa. Cosynergistic signaling by Fc*γ*RIIIa-pSyk and TLR9 enhances IL-17A and IL-21 production in Bcl6^+^ CD4^+^ T cells. Third, we show that Fc*γ*RIIIa-pSyk cosignaling differentially upregulates miRNA that contribute to T cell differentiation and TLR responses. Abundant IC formation and T_FH_ cells contribute to the autoimmune pathologies and infections [5, 38].

Increased frequency of T_FH_ cells are observed in the peripheral blood of SLE patients [6, 21, 37]. T_FH_ cell numbers correlate with an increase in autoantibody titers and tissue damage [39]. In addition to a strong TCR signal, CD28 cosignaling is required for the T_FH_ development [40]. However, the blockade of CD28 cosignaling in later stages of differentiation does not block the generation of T_FH_ cells [41]. This suggests the role of alternate activating co-signals in T_FH_ differentiation. In autoimmunity, signals other than CD28 drive the development of effector CD4^+^ T cells [42]. Our results suggest that the Fc*γ*RIIIa-pSyk is one of this unrecognized cosignal that can substitute the CD28 cosignaling in the T_FH_ development. *In vivo* Fc*γ*RIIIa^+^-pSyk^+^ CD4^+^ T cells were identified, which produce IL-21 and express Bcl6. Bcl6^+^ cells that produce IL-21, a T_FH_ cytokine show pSyk both *in vivo* and *in vitro*. Bcl6 expression has been reported in the circulating blood T_FH_ cells in SLE [6, 21]. These findings are contrary to previous report, where in circulating T_FH_ cells during HIV infection did not express Bcl6 [5]. It is proposed that low or lack of Bcl6 in circulating T_FH_ is a marker of the quiescent memory cells. It is somewhat surprising that Bcl6, a marker for GC T_FH_ cells is absent from circulating T_FH_ cells in viral infection, where one would expect activated cells. Bcl6 is necessary for GC reaction. The circulating T_FH_ cells should have the capacity to develop ectoid GCs, which are observed in the nephritic kidney and arthritic synovium. We also observed two populations of PD1 expressing cells, where pSyk^+^ cells produced lower amount of PD1. We are not clear, how pSyk participate in downregulating the PD1. We hypothesize that the Fc*γ*RIIIa-pSyk cosignal by downregulating PD1 favors cellular activation. PD1 targeted therapies provide significant clinical benefit in some patients, while temporary, partial or no clinical benefit in other patients [43].

Our *in vitro* data show that the Fc*γ*RIIIa-pSyk cosignaling in a milieu of four-cytokines IL1-β, IL-6, IL-23, and TGF1-β, results in the development of T_FH_ like cells [44]. These cells produce IFN-*γ*, IL-17A and IL-21 [17, 44]. In humans, IL-23 and TGF1β drives the early differentiation of T_FH_ cells, and cytokines IL-6, IL-21, and IL-1β support this process. Fc*γ*RIIIa costimulation upregulates GM-CSF and Csf2 RNA transcripts in naïve CD4^+^ T cells, genes normally associated with terminally differentiated T_H_17 pathogenic phenotype [17]. It is ambiguous which T_FH_ cell type is important for B cell help. One report showed that the T_FH_17 are efficient B cell helper for isotype class switching [1]. Another report suggested a role for blood T_FH_1 cells in SLE pathology [24]. Our results suggest a role of Fc*γ*RIIIa costimulation in the development of pathogenic CD4^+^ T cells and further authentication of T_FH_ subtypes in SLE is needed for therapeutic intervention. Excessive production of IL-21 from Fc*γ*RIIIa-pSyk cosignaling in T_FH_ cells could override their inhibition by the T_FR_ cells [27].

Our results also suggest a role for cosynergistic signaling from Fc*γ*RIIIa-pSyk and TLR9 in T_FH_ expansion (figure 4). A cosynergistic signaling between FcRs and NA-TLRs is observed in B cells, where DNA/RNA ICs engage BCR/FcRs and/or TLRs [29]. A coordinated signaling from BCR and TLR contributes to the positive selection of self-reactive B cells [45]. Chromatin-IgG complexes activate B cells by the dual engagement of IgM and TLRs [46]. In CD4^+^ T cells, DNA/RNA containing ICs by engaging Fc*γ*RIIIa and TLR9 could thus enhance T_FH_ development [18]. CpG 2006 bound to TLR9 on T cell surface and MyD88 protein co-immunoprecipitates with Fc*γ*RIIIa protein, which suggest a coordinated signaling in CD4^+^ T cells [18]. In B cells, joint signaling from ITAM (in FcRs) and MyD88 ( in NA-TLRs) is distinct, compared to individual signaling from either ITAM or MyD88 [29].

ICs are composed of large number of antigenic peptides [47]. Both FDC and/or GC B cells retain ICs on the cell surface, which fulfill the antigenic requirement for T_FH_ development in GC reaction. Our data suggest that the ICs bound to T_FH_ cells could achieve two objectives, first they could contribute to the fulfillment of excessive antigenic requirement of T_FH_ cells, and second they may provide an additional positive costimulatory signal. The additive positive costimulation by Fc*γ*RIIIa will contribute to TCR signal strength [16, 18]. A strong TCR signal is a prerequisite for the development of T_FH_ cells [40]. Fc*γ*RIIIa engagement by the ICs phosphorylates TCR signaling proteins [13, 16]. Mice studies have shown that the T_FH_ response is dictated by the amount of antigen, which correlate with the magnitude of GC B cell response [48]. The antigen presentation by GC B cells, and not the unique B cell derived signal is required for the development of T_FH_ cells [7]. A role for ICs in GC dynamics is also supported by the abundance of SCS macrophages in GCs, which retain and recycle ICs to cell surface and have poor phagocytic function [49]. Our *in vitro* experiments mimic some of these interactions and confirm a role for ICs in T_FH_ generation.

RNA-seq data revealed that Fc*γ*RIIIa engagement by ICs influence the expression of several key transcription factors required for T_FH_ development (figure 5). We observed the overexpression of GPCR signaling, FOXO and c-MAF pathway transcripts. Transcriptional regulators of FOXO and c-MAF pathways modulates T_FH_ responses [4]. PI3K signals via Akt promote T_FH_ differentiation. PI3-Akt signal phosphorylates FOXO, which leads to its exportation and degradation. This removes Bcl6 repression, thus promoting T_FH_ generation [50]. ICOS expressing cells are enhanced in SLE and these patients show more severe disease activity [37].

The miRNAs play a role in the development of CD4^+^ T_E_ cells. Our data provide an evidence that the Fc*γ*RIIIa mediated costimulation influences the CD4^+^ T cell responses by modulating the expression of miRNAs (figure 5). The miR-17~92 cluster, miR-146a, and miR-155 regulate T_FH_ development [4, 51]. Target genes of miR-17~92 have been identified to play an important role in the T_FH_ biology and IFN-*γ* production [52]. This cluster 17-92 is a master switch, which participate in T_H_1 and T_H_17 development [32]. The miR-155 and miR-146a have opposing roles in controlling T_FH_ differentiation [35]. miR-155 also enhances GC B cell differentiation and cytokine production. Several of these miRNAs regulate ICOS mediated responses. Upregulation of miR-1307 from Fc*γ*RIIIa-pSyk cosignaling suggest a mechanistic role for Fc*γ*RIIIa costimulation in SLE pathology via miR-1307 regulation [36]. Further validation of this data will be useful in recognizing events that contribute to autoimmunity.

Molecular mechanisms that provide stability to T_FH-_B cell cytoconjugates are critical in generating autoreactive B cells. Previous literature and our recent work have now confirmed the presence of low affinity FcRs on activated CD4^+^ T-cells [16, 18, 53, 54]. This opens up the possibility that the FcRs engagement by ICs could facilitate cytoconjugation, triggering the formation of lymphoid follicles observed in autoimmune pathologies. Both the timing and strength of the interactions between T_FH_ and B cells regulate GC dynamics. In GCs, B cells demonstrate longer interaction with T_FH_ cells compared to FDCs [55]. IC mediated interactions will be of several magnitudes higher in affinity, due to multiple bond formation, compared to other known protein pairs that hold T_FH-_B cells [10]. Furthermore, Fc*γ*RIIIa-pSyk cosignaling induces expression of IL21, PD1, ICOS and CXCR5, paired with their ligands they will further enhance T_FH_-B cell interactions. ICs bound to T_FH_ cell surface could likely engage receptors such as FcμR, Fc*γ*RIIb, Fc*γ*RIIc and BCR on B cells [56]. ICs mediated interactions could also be important for bidirectional signaling between the T and B cells during autoimmune pathology.

In summary, we provide evidence for a role of Fc*γ*RIIIa-pSyk signal in driving the expansion of T_FH_ cells. Furthermore, our data provide an evidence for a role of joint signaling by Fc*γ*RIIIa-pSyk and TLR9 in enhancing BCL6^+^ T_FH_ cells that produce IL-21. Fc*γ*RIIIa costimulation generates both blood T_FH_1 and T_FH_17 cells in *in vitro* from plate-bound ICs. Our data raises the possibility that Fc*γ*RIIIa^+^T_FH_ cells could be superior in providing B cell help due to their stronger association. Further exploring the role of low affinity FcRs in CD4^+^ T_FH_ cell responses will address several key questions that will enhance our understanding of the autoimmune pathology. Even though characterization of various T_FH_ subsets in viral pathology has been attempted, such characterization in SLE is still required. Further, understanding of ICs mediated interaction between T and B cells will be important to delineate autoimmune responses. A role of Fc*γ*RIIIa cosignaling in T_FH_ mediated responses should be further evaluated.

## Acknowledgments

Authors would like to thank Prof. Terry L. Moore for providing samples for the study. We would also like to thank the people from flow core for analyzing the samples. We would like to thank Drs. Kathie Mihindukulasuriya and Dale Dorsett from the Genomic Core at Saint Louis University.

